# DNA-encoded immunoassay in picoliter drops: a minimal cell-free approach

**DOI:** 10.1101/2022.09.20.508493

**Authors:** Barbara Jacková, Guillaume Mottet, Sergii Rudiuk, Mathieu Morel, Damien Baigl

## Abstract

Based on the remarkably specific antibody-antigen interaction, immunoassays have emerged as indispensable bioanalytical tools for both fundamental research and biomedical applications but necessitate long preliminary steps for the selection, production and purification of the antibody(ies) to be used. Here, we adopt a paradigm shift exploring the concept of creating a rapid and purification-free assay where the antibody is replaced by its coding DNA as a starting material, while exploiting a drop microfluidic format to dramatically decrease sample volume and accelerate throughput and sorting capability. The methodology consists in the co-encapsulation of a DNA coding for the variable domain of the heavy chain of heavy-chain only antibodies (VHH), a reconstituted cell-free expression medium, the target antigen and a capture scaffold where VHH:antigen accumulate to create a detectable signal, inside picoliter drop compartments. We first demonstrate successful synthesis of a functional hemagglutinin (HA)-tagged anti-GFP VHH, referred to as NanoGFP, at a high yield (15.3 ± 2.0 µg·mL^-1^) in bulk and in less than 3 h using PURExpress cell-free expression medium. We then use a microfluidic device to generate stable water-in-oil drops (30 pL) encapsulating NanoGFP-coding DNA, PURExpress medium, EGFP antigen and HA tag-specific magnetic nanoparticles prior to incubating at 37 °C the resulting emulsion under a magnetic field, inducing both *in situ* synthesis of NanoGFP and accumulation of NanoGFP:EGFP complexes on magnetically assembled particles. This allows us to assess, for the first time and in less than 3 hours, the binding of an antigen to a cell-free synthesized antibody, in a large number of picoliter drops down to a DNA concentration as low as 12 plasmids per drop. We also show that the drops of this immunoassay can be further sequentially analyzed at high throughput (500 Hz), thus offering capability for library screening, sorting and/or rare event detection. We finally demonstrate the versatility of this method by using DNA coding for different VHH (e.g., anti-mCherry protein), by characterizing VHH specificity in the presence of antigen mixtures, and by showing that antigens can be either inherently fluorescent or not. We thus anticipate that the ultraminiaturized format (pL), rapidity (3 h), programmability (DNA-encoded approach) and versatility of this novel immunoassay concept will constitute valuable assets for faster discovery, better understanding and/or expanded applications of antibodies.

## Introduction

Immunoassays are ubiquitous bioanalytical techniques, in which the presence of a target molecule is detected or quantified using antibody-antigen binding interaction.^[1]^ Despite tremendous importance in both fundamental research and real-world applications ranging from antibody discovery and analyte detection to medical diagnosis,^[2]^ their development remains time-consuming, costly and require dedicated human and instrumental resources. They necessitate the use of particular antibodies with proper affinity and selectivity against desired targets, resulting in long steps of antibody screening, production, purification and functionality characterization.^[3]^ Accelerating and simplifying such important biomolecular technique thus appears as a particularly timely challenge. Microfluidics and lab-on-chip technologies have the capability to dramatically decrease volume and time of reactions while being biocompatible, versatile and cost-effective.^[4]^ There have thus been many efforts in the past decade to develop robust microfluidic approaches to implement antibody bioassays in highly miniaturized formats. Drop microfluidics, which consists in handling, analyzing and sorting chemical and/or biological components in nano to picoliter monodisperse drops in a high-throughput fashion, has been identified as one of the most efficient methods to reach this objective.^[5,6]^ For instance, development of a microfluidic platform for compartmentalization, analysis and subsequent sorting of individual cells^[7]^ made possible to conduct studies on immune cells’ antibody secretome that was unexplorable by conventional flow cytometry. This breakthrough enabled not only better understanding of antibody secretion dynamics^[8]^ but also the characterization of antibody binding properties,^[8,9]^ both of which have facilitated the discovery of antibodies with desired functionality. These achievements had in common to be based on single-cell encapsulation in drops, with a particular focus on optimizing the methodology to increase the throughput and sorting capability of the developed microfluidic devices. Handling living cells was also accompanied by intrinsic constrains to maintain cells alive, such as limited time of experiments, mild conditions and use of biocompatible reagents. As an interesting complementary approach, and to further accelerate, diversify and simplify the capability of microfluidic antibody bioassays, we could think of substituting the confined secreting cell by a minimal and well-defined expression machinery producing the antibody of interest. So-called cell-free gene expression systems, in which a protein can be synthesized from coding DNA in a few hours, have been successfully exploited to synthesize functional proteins within a wide range of systems and applications.^[10,11]^ Beside their versatility and commercial availability, they are advantageously compatible with cytotoxic protein synthesis, artificial amino acid incorporations^[12,13]^ and new methods of extrinsic expression regulation, such as dynamic photocontrol.^[14]^ By simply using DNA coding for desired sequence, many protein types have already been synthesized including enzymes,^[15–17]^ membrane proteins^[18–21]^ or large protein assemblies.^[22–26]^ Interestingly, cell-free expression systems can also be employed in miniaturized format, successful examples including fluorescent proteins,^[27–29]^ enzymes^[30–33]^ and transcriptional regulators.^[34]^ In contrast, due to the large size and complex higher structure of immunoglobulins (IgGs), their synthesis has so far only been achieved in bulk cell-free systems, after substantial efforts to optimize both antibody-coding DNA sequence and the composition of cell-free expression medium.^[35–37]^ To achieve functional antibody synthesis in minimal compartments without optimization steps, we could suggest instead to synthesize the variable domains of the heavy chain of heavy-chain only antibodies (VHH)^[38]^ engineered from naturally occurring antibodies found in camelids. VHH are small-sized (∼13 kDa), stable, single-unit and easily-foldable proteins, originally used as tools for intracellular protein tracking^[39]^ or reagents for high-affinity protein purification.^[40]^ They nowadays represent a highly promising alternative to conventional therapeutic antibodies.^[41,42]^ To our knowledge, cell-free synthesis of VHH has up to now only been reported in a bulk format, in particular for antibody discovery purposes, using techniques of mRNA, ribosome and phage displays.^[43–45]^ Here, we propose not only to express functional VHH in microfluidic drops by cell-free expression from its coding DNA for the first time, but also to concomitantly, and in the same drops, assess the capability of the synthesized antibody to selectively capture its target antigen. We first synthesized in bulk an anti-green fluorescent protein (GFP) VHH, referred to as NanoGFP, using a reconstituted cell-free expression system and characterized both the synthesis yield and the functionality of the synthetized VHH. By co-encapsulating coding DNA, expression machinery, a capture scaffold and EGFP antigen in microfluidic-generated picoliter drops, we assessed the antibody-antigen interaction along the course of expression (a few hours) inside individual drops. Finally, using a laser detection system, we determined, at a high-frequency and in a large number of individual drops, the performance of this picoliter immunoassay in terms of minimum number of DNA copies per drop, antigen detection limit and capture selectivity.

## Results and discussion

Figure 1 depicts the central concept of our approach. It consists in encapsulating in the same picoliter compartment, the minimal components necessary to both synthesize a desired VHH and characterize *in situ* its binding affinity for a target antigen. Water-in-oil drops are used to encapsulate a small number of DNA coding for the VHH, a cell-free expression medium to synthesize the VHH from DNA and specific and/or dummy antigen(s). To detect the VHH-antigen binding, the strategy consists in implementing a capture scaffold onto which VHH are tethered upon their cell-free synthesis. As a result, target antigens, initially homogenously distributed inside the drop content, accumulate on the VHH-decorated capture scaffold, resulting, in the case of labelled VHH and/or antigens in a signal accumulation from the antibodies and/or its bound antigens. This concept combines several advantages. No cell is involved, avoiding all steps of cell maintenance, and expanding the range of conditions that can be explored. All components in the drops are supplied in a known and prescribed concentration, offering reliability and robustness. The duration of the whole assay, starting from the DNA encapsulation to the detection of the antigen-binding, is mainly determined by the cell-free expression reaction rate, and is thus of the order of a few hours only. Finally, using VHH-coding DNA as starting material enables easy applicability to virtually any kind of VHH sequence while the micrometric drop format offers immediate compatibility with microfluidic handling, such as high-throughput drop production, testing and sorting. To demonstrate this concept, we focused on a few important key-steps: picoliter drop production and encapsulation of the minimal assay, *in situ* VHH synthesis and concomitant antigen-binding analysis, and high-frequency analysis in a large number of flowing drops. As a model antigen, we mainly used enhanced green fluorescent protein (EGFP), a commonly used and fluorescent target. We associated it to an anti-GFP VHH, referred here to as NanoGFP^[46]^ which was encoded in the encapsulated DNA. For the cell-free VHH synthesis, we focused on reconstituted PURE expression system,^[47]^ a minimal set of recombinant components purified from *E. coli*, for their well-known composition, fast protein synthesis and commercial availability.

**Figure 1.**
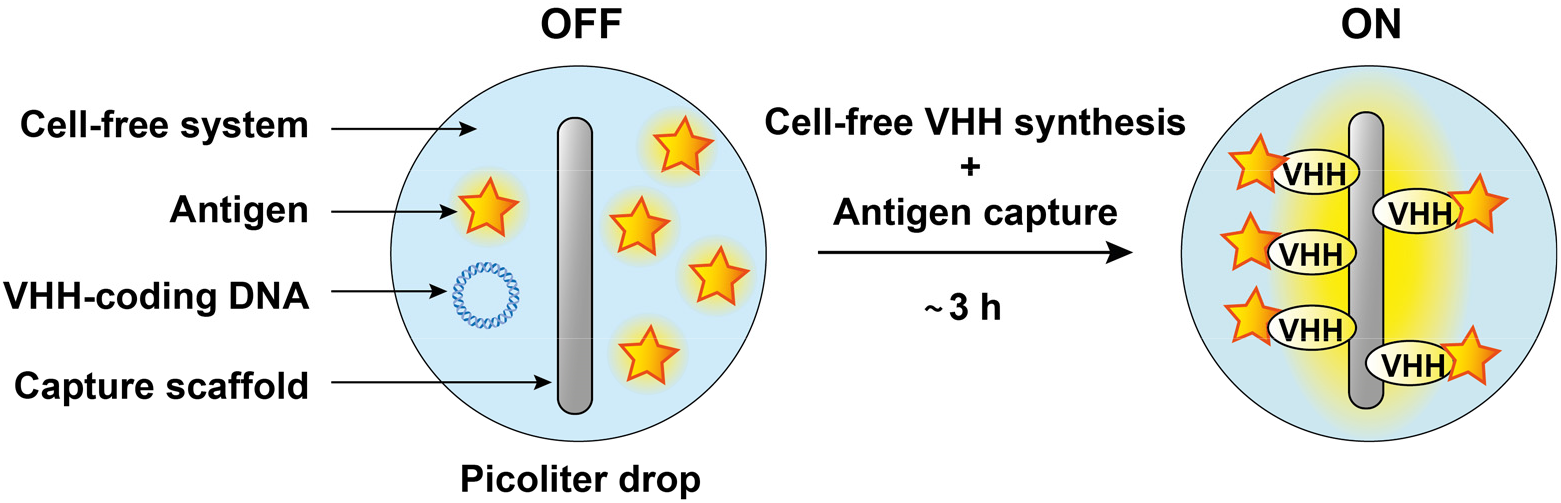
Cell-free DNA-encoded immunoassay in picoliter drops allows rapid assessment of antibody-antigen binding with minimal components, programmability and DNA instead of purified antibody as starting material. The concept is based on water-in-oil drops of micrometric size containing cell-free expression mix, fluorescent antigen, VHH-coding DNA and a capture scaffold capable to attach the antibody (VHH) upon the synthesis. The resulting emulsion is incubated at 37°C for 3 h during which the VHH is cell-free synthesized, is tethered onto the capture scaffold and binds the antigen in case of sufficient antibody-antigen affinity. This binding alters the distribution of antigen within the drop, from unbound antigen producing a homogeneous fluorescent signal (OFF) to antigen accumulated on the capture scaffold, resulting in a local increase in fluorescence intensity (ON).

Prior to drop encapsulation, we characterized the amount of NanoGFP that could be synthesized by PURExpress® in bulk and further tested its antigen recognition capability. To this end, we designed a plasmid containing the necessary elements for transcription and translation in PURE system (T7 promoter, ribosome binding site (RBS) and T7 terminator) around the gene coding for VHH and hemagglutinin (HA) tag separated by a linker region (Fig. 2A). For the VHH, we used a sequence previously optimized for expression in *E. coli*.^[46]^ The HA tag, designed for tethering the synthesized protein onto the signal amplification scaffold, was positioned in the C-terminus to minimize any possible effect on the antigen binding at the VHH paratope region^[48]^ This fusion increased the protein molecular weight by 2 kDa only, leading to an overall size of 16.4 kDa. The resulting plasmid was incorporated into a conventional PURExpress® mix and incubated at 37 °C for 3 hours in a tube. Capillary western blot analysis of the product revealed a sharp single band, showing that the synthesized NanoGFP was properly produced and as full-length monomers only (Fig. 2B). Its position was slightly above that obtained with a commercially available anti-GFP VHH of 13.9 kDa, in agreement with its expected size. ELISA titration revealed a yield of 15.3 ± 2.0 µg·mL^-1^ of synthesized NanoGFP in PURExpress® (Fig. S1). Replacing PURExpress® by PURE*frex*®, another PURE cell-free expression medium, resulted in successful yet lower-yield NanoGFP synthesis. Furthermore, adding supplements such as disulfide bond enhancer or chaperones did not improve the yield (Fig. S1), showing the interest of working with short-sized and simply structured proteins such as VHH. We next confirmed the binding activity of the cell-free produced VHH. To this end we follow the binding of EGFP as a function of NanoGFP concentration to establish the dose-response curve (Fig. 2C). The method was validated with a commercially available VHH (unknown complementarity determining region – CDR – sequences) leading to an apparent dissociation constant *K*_*D*_^*app*^ = 0.49 ± 0.03 nM. Interestingly the cell-free expressed NanoGFP led to a similar value *K*_*D*_^*app*^ = 0.26 ± 0.03 nM, which was also in the same range as what was reported with the same VHH produced in bacteria.^[39,49]^ All these results show that cell-free synthesis in PURExpress® system produced functional NanoGFP at conventional yield and expected binding affinity. We next exploited the DNA-encoding approach of our strategy and applied it to explore the cell-free synthesis of other VHH sequences. We started with several anti-GFP variants known for their different binding affinities and showing significant sequence diversity especially in the CDR3 region (Fig. S2). Interestingly, all synthesized VHH also displayed a single and well-defined band in Western analysis and with ELISA titration ranging from 2.6 ± 0.1 to 60.7 ± 10.0 µg·mL^-1^, with PURE*frex*® leading to similar results with lower yields of expression (Fig. S3). Dose-response analysis in the accessible concentration range showed functional binding activity of the synthesized mutants (Fig. S4). Using a cell-free expression medium from *E. coli* purified components was thus found to be a valuable strategy to synthesize significant amounts of functional anti-GFP VHH in bulk, in one step and in a few hours only, while allowing to explore various VHH sequences by simply modifying its encoding DNA.

**Figure 2.**
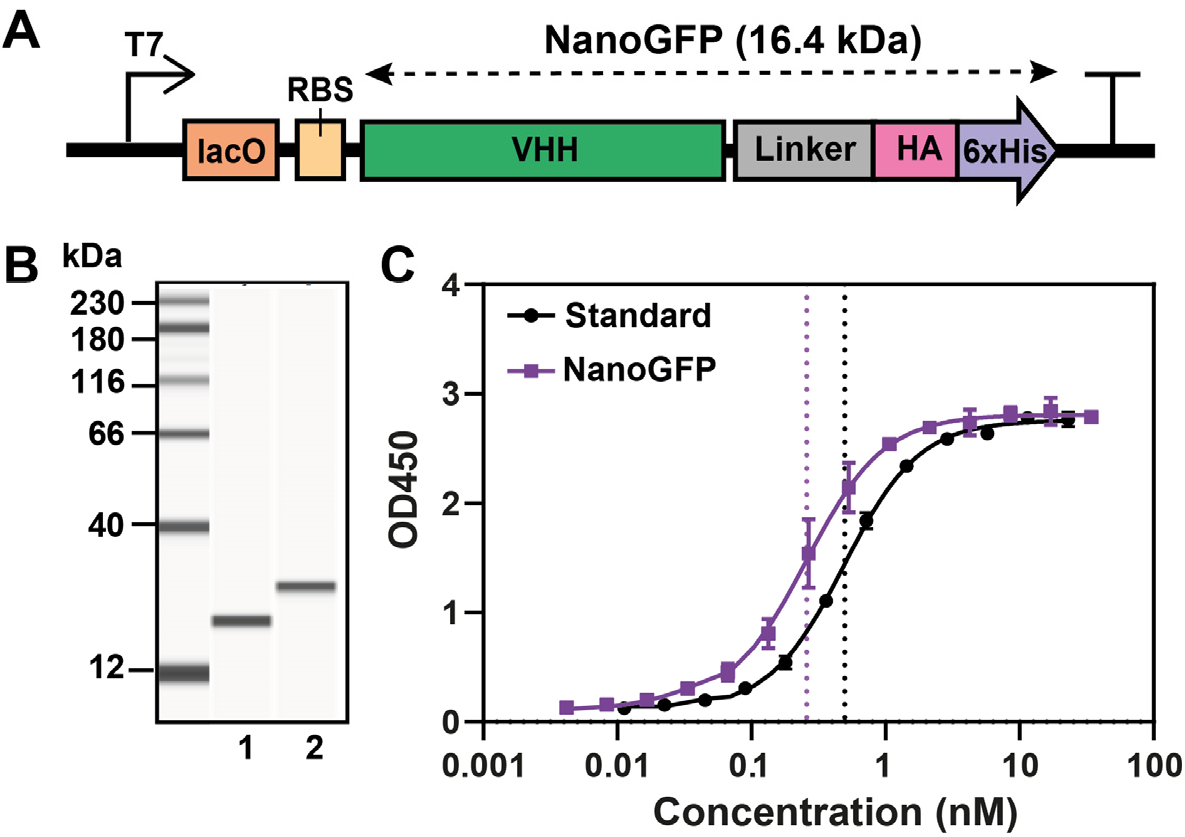
Characterization of anti-GFP VHH, called NanoGFP, expressed by PURExpress® in bulk format. A) Schematic representation of the DNA template used for NanoGFP cell-free expression. The template contains T7 promoter, lac operator (lacO) and ribosome binding site (RBS) located upstream of the gene coding for NanoGFP (16.4 kDa), composed of VHH-coding gene separated by a linker from HA epitope tag and 6xHistidine tag, followed by a T7 terminator. B) Capillary western blot analysis of commercially available anti-GFP VHH (lane 1) and cell-free expressed NanoGFP (lane 2). C) Dose-response curves of commercially available anti-GFP VHH (Standard) and cell-free expressed NanoGFP. The curves were obtained by indirect ELISA, using EGFP antigen for VHH capture and an anti-VHH peroxidase-conjugated IgG for detection. Dotted lines indicate the apparent dissociation constant, for the standard: *K*_*D*_^*app*^ = 0.49 ± 0.03 nM and for the NanoGFP: *K*_*D*_^*app*^ = 0.26 ± 0.03 nM. NanoGFP was expressed at [DNA] = 4 ng·µL^-1^, 37°C, 3 h.

Experiments described before involved reaction volumes of about 25 µL and a large number of DNA copies per reaction (10^10^). To establish a DNA-encoded immunoassay in 10^6^-time smaller volumes (Fig. 1), we had to devise a method to confine a small number of VHH-coding DNA molecules in highly miniaturized containers and implement a method allowing to detect *in situ* the functionality of the synthesized VHH. The strategy was to use a microfluidic device to produce picoliter drops co-encapsulating VHH-coding DNA, the cell-free expression medium, the EGFP antigen and a capture scaffold allowing to analyze the binding of the antigen to the VHH synthesized *in situ* (Fig. 3). This binding was assessed using a capture nanoparticle made of a streptavidin-coated superparamagnetic bead previously functionalized with a biotinylated IgG specific to the HA tag in the C-terminus of the target synthesized VHH (Fig. 3A, *left*). The capture nanoparticles were assembled together with the VHH-coding DNA and the EGFP antigen. This mix and PURExpress® expression medium were injected at 4 °C as the two aqueous phases co-flowing in a microfluidic-device where drops were generated at a flow-focusing junction with a fluorinated oil supplemented with fluorinated surfactants to ensure non-coalescence of the produced drops^[50]^ (Fig. 3A, *middle*). The advantage of using capture particles of nanometric dimensions (∼ 300 nm in diameter) was to enable working at a concentration high enough to avoid heterogenous distribution (Poisson partitioning) while offering a large surface area for binding. Our device typically produced 2.5·10^6^ drops of 30 ± 3 pL of narrow polydispersity in 10 min. The resulting emulsion was collected and immediately incubated at 37 °C under application of a magnetic field resulting in the formation of a bar of aligned particles^[8]^ forming a capture scaffold in each drop (Fig. 3A, *right*). Microscopic observation on a large number of individual drops in parallel revealed that EGFP signal was increasing at the position of the assembled capture particles, while vanishing in the background (Fig. 3B, S5). The characteristic diffusion time for a NanoGFP:EGFP complex of 44.4 kDa was estimated to be 4.5 s (Text S1) using the drop diameter as a characteristic size, allowing us to follow, with good temporal resolution, the capture of antigen along the course of VHH expression (Fig. S6A) which is known to be sustained for about 3 h in bulk (Fig. S7). The same experiment performed without DNA resulted in no signal evolution (Fig. S6B), demonstrating that the increase of EGFP fluorescence in the presence of the coding DNA was resulting from accumulation of synthesized VHH binding the antigen. To our knowledge, this constitutes the first *in situ* observation of antibody-antigen binding in a minimal cell-free system at picoliter scale. To compare the antigen binding signal evolution to the actual synthesis of VHH, we determined in each drop the average signal intensity at the particle position and established its distribution among 300 individual drops as a function of time. We measured independently the amount of synthesized VHH at each time point using ELISA after breaking the emulsion. We found that the signal from EGFP accumulated at the capture scaffold was indeed correlated with the evolution of VHH level along the course of its expression (Fig. 3C). Interestingly, significant signal concentration could already be observed in less than 30 min. After 175 min of incubation at 37 °C, a concentration of 6.7 nM of synthesized VHH resulted in a 5-fold increase of the EGFP signal per drop in average. Note that this assay involved a number of DNA copies per drop λ = 300. Cell-free expression at the same DNA concentration in bulk led to a yield of 4.9 nM of synthesized VHH after the same incubation time, emphasizing that the encapsulation strategy did not hamper and may even favor the *in situ* protein synthesis. All these results show that squeezing anti-GFP VHH-coding DNA, its target antigen and a capture scaffold in microfluidic-generated picoliter drops allows one to quickly achieve VHH synthesis and concomitantly assess its functional binding.

**Figure 3.**
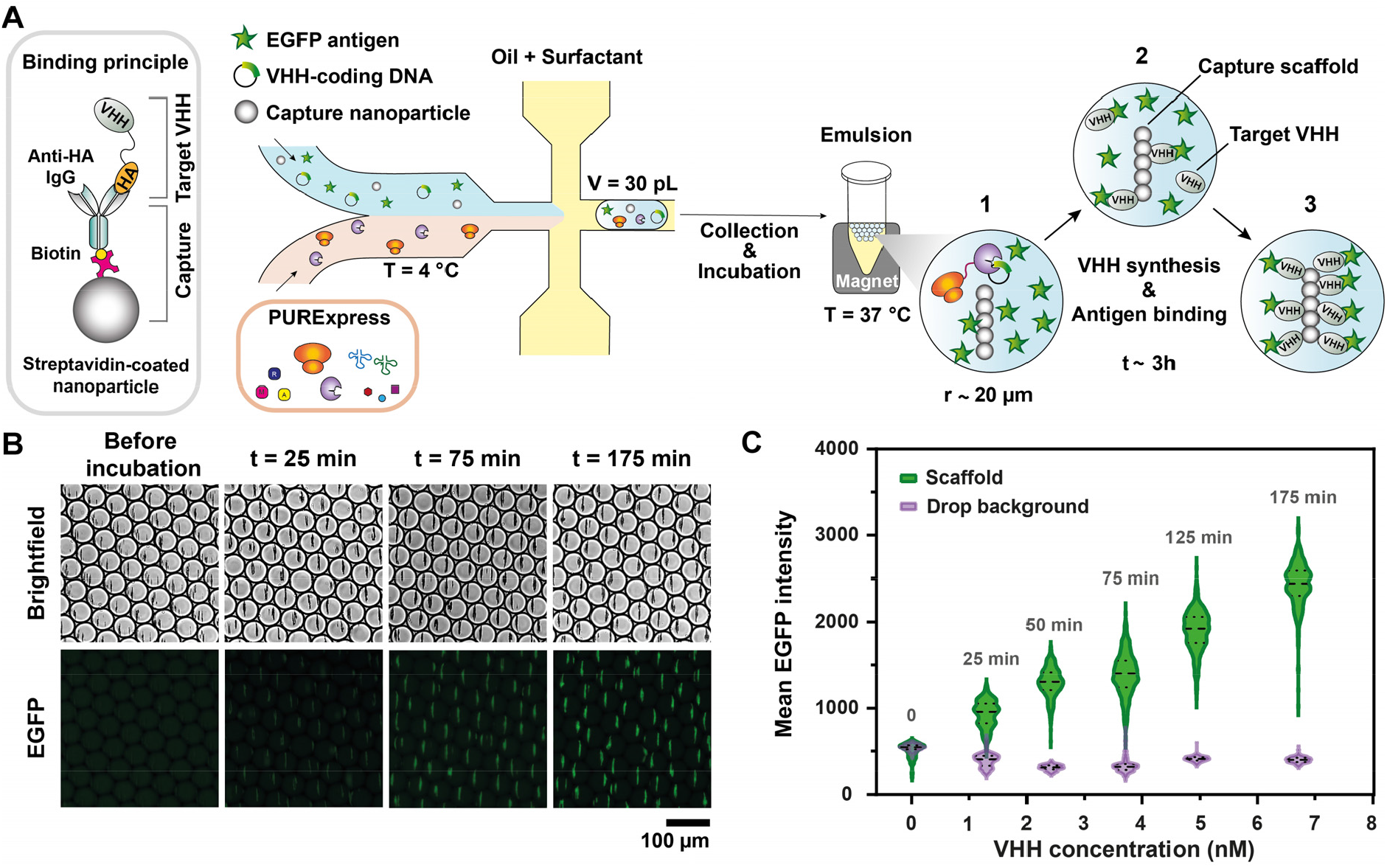
DNA-encoded immunoassay produced by microfluidics and performed with DNA coding for NanoGFP, allows for real-time *in situ* observation of VHH:EGFP binding thanks to accumulation of the complex on the capture scaffold. A) Microfluidic workflow for immunoassay preparation and the mechanism of formation of fluorescent readout. Streptavidin-coated magnetic nanoparticles (300 nm in diameter) were functionalized with biotin-conjugated anti-HA IgG to allow the capture of target VHH and suspended in a solution containing VHH-coding DNA (λ = 300 plasmids per drop) and EGFP (40 nM). The mix was injected in a droplet generator with a flow-focusing geometry, in a co-flow with PURExpress® (400 µL·h^-1^) and emulsified by fluorinated oil with 2% fluorosurfactant (1400 µL·h^-1^), resulting into monodisperse drops of ∼ 30 pL. The collected emulsion was incubated at 37 °C under a magnetic field to align the magnetic nanoparticles and during 3 h the VHH was progressively synthesized (1) and tethered on the capture scaffold together with its bound EGFP (2) which produced a bright and localized fluorescence signal (3). B) VHH expression and antigen binding followed by fluorescence microscopy. After encapsulation, the emulsion was injected into a glass microfluidic chamber and imaged before and during incubation at 37 °C. C) Violin plot of the mean EGFP intensity distribution measured on the scaffold and in the drop background as a function of synthesized VHH concentration measured by sandwich ELISA on a broken emulsion. Mean EGFP intensity of 300 droplets was assessed by particle analysis on background-subtracted images (dashed line: median, dotted lines: lower and upper quartiles). The time of incubation is indicated above each distribution.

After characterizing a large ensemble of drops in parallel, we sought after sequential analysis of individual drops at a high frequency offering, for instance, the capability to detect rare events in real time. This method was implemented not only to characterize the performance of the assay but also to devise a method that could be readily compatible with *in situ* sorting. To this end, we injected the cell-free expression emulsion after 3 h incubation into another microfluidic device integrating side oil channels to separate the drops and a laser-assisted *in situ* fluorescence measurement (Fig. 4A, *left*). In this system, the whole width of the channel was laser-illuminated to ensure uniform profile of each drop flowing through the measurement window and the fluorescence emission was recorded by a photomultiplier tube (PMT) detector. The detection of the maximum intensity in each drop fluorescence profile, referred to as Max EGFP, allowed us to discriminate between situations such as no detectable VHH expression (e.g., due to absence of DNA or low-yield synthesis), expression of non-binding VHH (e.g., lack of affinity), and successful synthesis of functional VHH (Fig. 4A, *right*). In the first two cases, Max EGFP was in the range of the background EGFP signal, while the presence of the functional VHH was characterized by a Max EGFP value superior to the fluorescent background produced by unbound EGFP. Drops were analyzed at a typical frequency of 500 Hz, offering the possibility to explore a pool of thousands drops in a few seconds only. In particular, we analyzed the drop population produced under the conditions of Fig. 3 and analyzed the distribution of Max EGFP, with or without VHH-coding DNA (λ = 300, Fig. 4B). Without DNA, the distribution was very sharp and at low Max EGFP values, allowing us to define a threshold (dashed line) above which larger values of Max EGFP would indicate the presence of the produced VHH binding its fluorescent antigen (positive drops). Interestingly, with DNA the signal was more broadly distributed and the majority of drops (92 %) were above the binding threshold, demonstrating both successful synthesis of functional VHH as well as a good sensitivity of the detection method. In this assay, it was important to optimize the EGFP concentration as the performance of the detection resulted from a tradeoff between antigen amount sufficient to ensure binding detection and low enough to avoid a too strong background inside the drop. For this purpose, we determined how EGFP concentration at a fixed λ = 300 affected the distributions of Max EGFP values and we selected a concentration of 40 nM for which positive drops displayed particularly high signal compared to the background (Fig. S8). We then assessed the minimum number of DNA copies per drop that could be used at this EGFP concentration for the assay to remain applicable. We thus performed our analysis on different emulsions produced with a varying VHH-coding DNA concentration (Fig. 4C, Figs. S9). We found that positive drops could be detected at concentrations as low as λ = 12, the fraction of positive drops significantly increasing with an increase in λ. This correlates with the amount of VHH produced in drops. For instance, the VHH concentration for λ = 12 was measured to be 0.24 nM, which is about 30 times less than with λ =300. Notably, at such a low concentration, the fraction of NanoGFP (*K*_*D*_^*app*^ = 0.26 ± 0.03 nM) bound to its antigen can be estimated to be 48%, showing that the detection limit is a signature of inherent VHH affinity. To further demonstrate the applicability of this assay we added a secondary fluorescent antibody conjugate specific to GFP. By performing the same single-drop analysis on both fluorescent signals (Fig. S10) we found that 79% of drops presented secondary antibody signal above the threshold, when 89% of this population displayed EGFP signal above threshold. This shows the possibility of multiplexing the assay as well as a way to detect non fluorescent antigens. All these results demonstrate the capability of this method to sequentially analyze large numbers of single drops containing only several copies of VHH-coding DNA and determine the VHH functionality against virtually any kind of antigens, inherently fluorescent or not.

**Figure 4.**
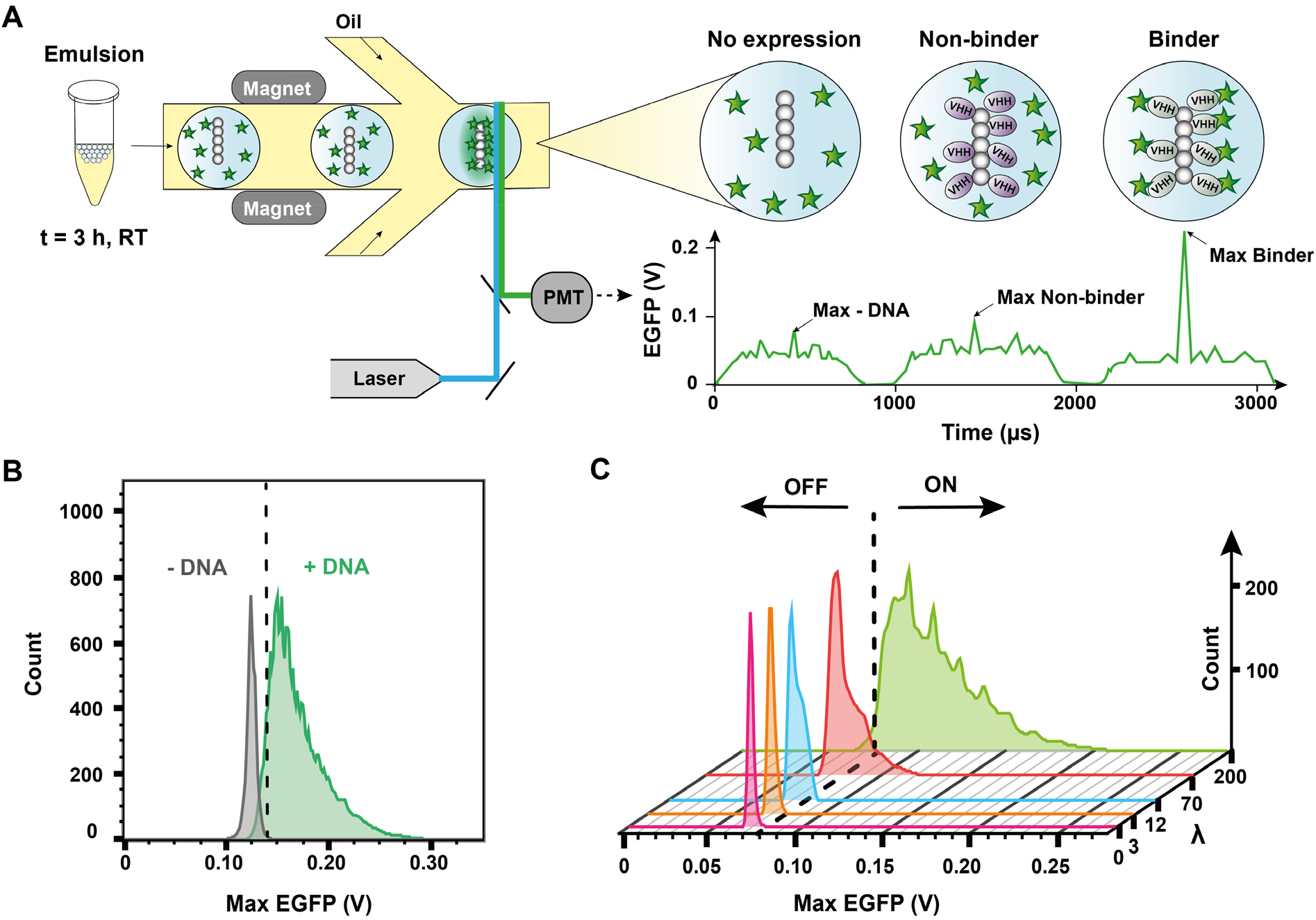
Sequential analysis of a large number of droplets reveals VHH-antigen binding can be detected with DNA concentrations as low as λ = 12 plasmids per drop. A) Laser and PMT-equipped microfluidic platform provides a means to differentiate droplets containing target antibodies (binder VHH) from the rest (not expressed or non-binder VHH). After 3h at 37°C the emulsion is injected at RT into droplet analysis device with magnets used to align the capture scaffolds. Drops are spaced by fluorinated oil with 0.5% of fluorosurfactant (f = 250 Hz), excited by a 488 nm laser. Emitted photons are converted by a PMT into an electrical signal EGFP (V), displayed as a function of time. Droplets containing binder VHHs showed a higher maximal EGFP intensity (Max EGFP) due to accumulation of EGFP on the linear capture scaffold and are defined as positive. B) Max EGFP distributions of negative control (-DNA, λ= 0) and emulsion with DNA coding for NanoGFP (+DNA, λ= 300). Highest Max EGFP detected for -DNA was fixed as the threshold above which droplets were considered as positive (dashed line). C) Max EGFP distributions for λ = 0, 3, 12, 70 and 200. All emulsions contained EGFP (40 nM). The exact numbers of analyzed droplets (Table S1) with fractions of positivity per condition (Fig. S9) are available in SI.

All previous experiments involved a single type of antigen-antibody interaction. To investigate the specificity of cell-free synthesized VHH, we performed the sequential analysis method in the presence of several antigens. As a proof of concept, we encapsulated simultaneously EGFP and the red fluorescent protein mCherry in the drops and we measured both green (EGFP) and red (mCherry) fluorescent signals in the presence of different VHH-coding DNA (Fig. 5). With DNA coding for NanoGFP, the majority of drops (85%) were positive in the green channel while all the drops showed negative signal in the red channel, meaning selective NanoGFP:EGFP binding (Fig. 5, *top*). Conversely, with the same amount of DNA (λ = 300) but coding for anti-mCherry VHH (LaM-4)^[51]^, positive drops were detected with red channel only (43%, Fig. 5, *middle*). The lower proportion of positive drops for red signal is likely attributed to the lower yield of expression of LaM-4 (2.8 ± 0.6 µg·mL^-1^ in bulk, Fig. S3), which is approximately 5 times less than that of NanoGFP, and the lower brightness of its mCherry antigen. Without DNA (Fig. 5, *bottom*) no positive signal was detected in neither of channels confirming that the positive peaks in the presence of DNA evidenced selective antigen binding.

**Figure 5.**
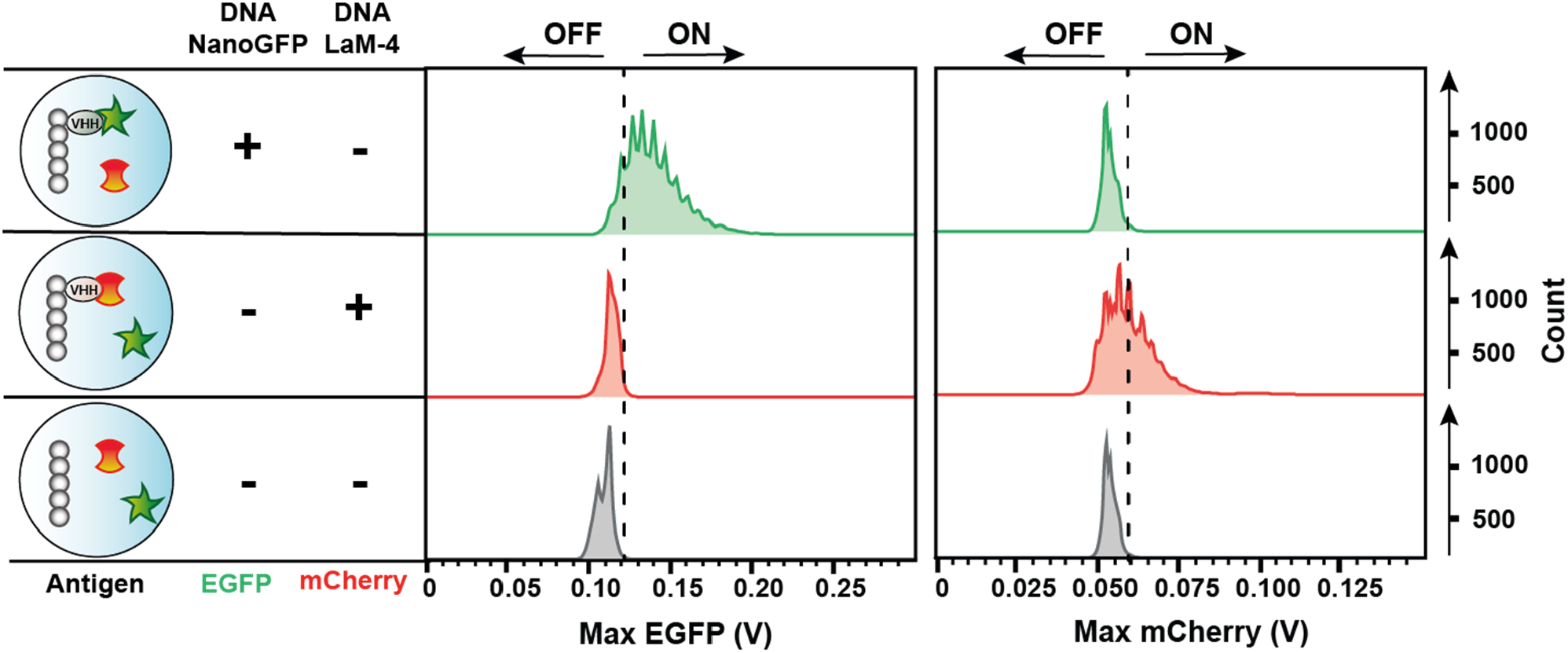
Cell-free expressed NanoGFP and LaM-4 present selective antigen binding in the presence of both their target antigens (EGFP and mCherry, respectively). Emulsions with DNA coding for NanoGFP (top row), DNA coding for LaM-4 (middle row), and without DNA (bottom row), all containing both EGFP (40 nM) and mCherry (80 nM), were incubated for 3 h at 37°C and analyzed using 488 nm and 561 nm lasers. The detection thresholds (dashed lines) for both channels were determined by the highest Max EGFP and Max mCherry detected in emulsion without DNA. The exact numbers of analyzed droplets are available in SI (Table S1).

## Conclusion

We have demonstrated the possibility to perform an immunoassay using a VHH-coding DNA as a starting material instead of a purified antibody as it is conventionally done. By removing all cell handling and purification steps, functional VHH was synthesized by cell-free expression and its capability to bind its specific antigen was directly characterized *in situ*. The whole process took a maximum of 3 hours, with binding detectable in less than 30 min, and required minimal amounts of materials and reagents. The assay was performed inside water-in-oil picoliter drops encapsulating the coding DNA, a cell-free expression medium with purified components from *E. coli* (PURExpress®, PUREfrex®), the target antigen, and magnetic particles accumulating on their surface the synthesized VHH binding its antigen. Simply incubating such microcompartments at 37 °C under a magnetic field allowed us to follow the accumulation of the antigen signal on the self-assembled nanoparticles and therefore assess for the first time the functionality of a cell-free expressed VHH binding its antigen. The hemagglutinin (HA)-tag was conveniently used as a generic and easy-to-implement way to link the synthesized VHH to the particles but other conjugation methods could be envisioned. Similarly, we chose magnetic nanoparticles for their large surface area of binding and capability to self-assemble under command once encapsulated to avoid Poisson partitioning, but other capture scaffolds could be used as well. The concept of the immunoassay was demonstrated here mainly using a DNA coding for an anti-GFP VHH and the corresponding EGFP antigen. Using DNA instead of a preliminarily purified antibody confers to the method a unique degree of programmability that was demonstrated with the successful cell-free synthesis and characterization of different mutants against the same antigen as well as VHH targeting other antigens (mCherry). By simply adapting the DNA sequence, the method thus offers not only the possibility to virtually implement any VHH but also to modify them in a highly tunable manner (e.g., tag addition, artificial amino acid incorporation, protein truncation/fusion). We also showed that the antigen detection was operational with antigens that can be inherently fluorescent (here, EGFP, mCherry) or not (secondary antibody labelling). For a fully DNA-encoded approach, we are also currently implementing the *in situ* cell-free expression of the antigen itself (data not shown). The panel of antibody-antigen interactions that can be explored with this method thus appears to be potentially extremely large. The microfluidic format of the assay (drop generating device) only requires standard device fabrication and set-ups, thus being implementable in a broad variety of environments, while offering the possibility to work with minimal amounts of reagents at a high speed. We have shown in particular that the picoliter drops containing the DNA-encoded immunoassay could be analyzed individually, in a parallel or sequential manner, at a high frequency (500 Hz) and with amounts of DNA as low as 12 copies per drop. In a synthetic biology context, this work reveals a new facet of cell-free expression that now enables the study of antibody-antigen interaction in a drastically simplified yet highly programmable format. From a biotechnological point of view, this study describes a methodological paradigm shift in immunoassay where genetically active synthetic microcompartments produce and directly report the binding activity of their encoded antibody, thus constituting a promising tool for faster discovery and improved implementation of antibodies.

## Supporting information

This file includes: Materials and methods; Supplementary Figures S1-S10; Supplementary Tables S1; Supplementary Text S1; Supplementary references

## Acknowledgement

We thank Nathalie Coulteaut (Sanofi) and Fréderic Lacroix (Sanofi) for providing access to equipment and experimental support; Vasily Shenshin (Sanofi) for assistance in microfluidic handling; Micaela Vitor (Sanofi), Hélène Erasimus (Sanofi) and Samy Dehissi (Sorbonne Université) for helpful scientific discussions.

