## Supplementary material for "DNA-encoded immunoassay in picoliter drops: a minimal cell-free approach": This file includes: Materials and methods; Supplementary Figures S1-S10; Supplementary Tables S1; Supplementary Text S1; Supplementary references

**Table of contents:**

1. Materials and methods
2. Supplementary Figures S1-S10
3. Supplementary Tables S1
4. Supplementary Text S1
5. Supplementary references

### 1. Materials and methods

##### VHH sequences and cloning into expression vector

Different sequences of anti-GFP and anti-mCherry VHHs were obtained from published works.[1,2] To minimize expression variability between sequences, their DNA coding sequences were redesigned to keep all framework regions constant and were synthetized by GeneArt (ThermoFisher Scientific) as linear fragments. They were subcloned into a plasmid derived from pRSET-B vector (ThermoFisher Scientific) using Gibson assembly master mix prepared according to previously published protocol[3].A representative plasmid map, the lists of sequences of interest and primers used for subcloning are available below.

##### DNA preparation

For each VHH sequence, plasmid was replicated in NEB ® 5-alpha competent *Escherichia coli* (New England Biolabs). After overnight culture (12-16h) in 4 mL of Luria Broth (LB) at 37°C and 220 rpm DNA was extracted using Monarch MiniPrep kit (NEB). DNA concentration was measured by Qubit 4 Fluorometer (ThermoFisher Scientific), using dsDNA Broad Range Assay and DNA purity was assessed by measurement of $A_{260}/A_{280}$ ratio on BioSpectrometer® (Eppendorf).

##### Cell-free protein synthesis

*In vitro* protein synthesis was carried out using PURE*frex*® (GeneFrontier) or PURExpress® (NEB) according to manufacturer's instructions. Due to the observed higher yield in bulk (Fig. S3), PURExpress was preferred for droplet-based expression. Unless mentioned otherwise, a concentration of 4 ng/µL of circular DNA (with typical size of 3-4 kb) was used for expression together with 0.8 U/µL of RNAse Inhibitor, Murine (NEB) to minimize potential RNAse contaminations. To reach maximal expression, reaction mix was incubated for 3 h at 37°C. Reaction medium was either analyzed directly post-synthesis or stored at -20°C for no longer than 1 month.

##### Western Blot

Cell-free reaction medium was analyzed after 3 h of incubation using fully automated immunoblotting SimpleWestern™ platform Jess (ProteinSimple, Bio-Techne). Briefly, samples were diluted 1:5 in Antibody Diluent, part of Anti-Goat Detection Module (ProteinSimple, Bio-Techne), and processed according to manufacturer's instructions. 12-230 kDa separation module was then used to electrophoretically separate protein constituents based on their molecular weight. After immobilization within capillary, proteins synthesized in PURExpress® were immunoprobed with primary antibody AffiniPure Goat Anti-Alpaca IgG, VHH domain (Jackson ImmunoResearch) and those synthesized in PURE*frex*® with anti-His IgG (R&D systems) ,both at 1:50 dilution and quantified by chemiluminescence using the HRP-based Anti-Goat Detection module. Digital capillary image of chemiluminescent signal was captured and displayed in lane view and as an electrophoretogram using Compass for SimpleWestern software (Version 5.0.1, Protein Simple).

#### ELISAs

Concentrations of functional cell-free synthetized VHH, either from bulk or from droplet expression (see emulsion generation below), were measured by sandwich enzyme-linked immunosorbent assay (ELISA). To monitor dynamics of VHH expression in droplet compartment, fractions of emulsion were collected every 25 minutes, combined with 1H,1H,2H,2H-Perfluoro-1-octanol (97%, Sigma-Aldrich) in 1:1 ratio, thoroughly mixed and centrifuged to isolate aqueous phase. To stop protein production, ribosome inhibitor chloramphenicol (Sigma-Aldrich) was added to final concentration of 40 µg/ml.

To perform sandwich ELISA, 96-well plates with hydrophilic surface (MaxiSorp) were coated with AffiniPure Goat Anti-Alpaca IgG, VHH domain antibody at 2.5 µg/mL in coating buffer (16 mM $Na_2CO_3$, 34 mM $NaHCO_3$, pH=9.6) and incubated overnight at 4°C. Plates were then washed twice with wash buffer (phosphate buffered saline DPBS with 0.05% v/v Tween® 20 (MP Biomedicals)), and incubated 1 h at room temperature (RT) with blocking buffer (wash buffer with 1% w/v of bovine serum albumin (BSA, Sigma-Aldrich)) to prevent non-specific binding. After two additional washing steps, a part of wells was incubated with a range of concentrations of protein standard Alpaca anti-GFP VHH (ChromoTek) and the rest with serial dilutions of samples to quantify. All protein dilutions were realized in blocking buffer. After 1 h incubation at RT plates were washed 5 times and incubated with secondary horseradish peroxidase (HRP) conjugated AffiniPure Goat Anti-Alpaca IgG, VHH domain antibody (Jackson IR) at 1:10 000 dilution during 1h. Plates were washed 5 times and incubated for 5 min with 3,3',5,5'-Tetramethylbenzidine (TMB) to produce chromogenic readout. Enzymatic reaction was then blocked by addition of 1N hydrochloric acid (HCl) and optical density (OD) of samples was measured at 450 nm using a microplate reader (Infinite M Plex, Tecan). To estimate the concentration of analytes, samples data were interpolated to four parameter logistic curve obtained with data produced by dilutions of protein standard (Fig. S1). For samples synthetized in bulk, measurements were realized in triplicates and for those extracted from emulsion in duplicates.

The apparent dissociation constant (EC50) was estimated using indirect assay format. 96-well plates with hydrophilic surface (MaxiSorp) were coated with Enhanced Green Fluorescent Protein (EGFP, ChromoTek) at 2 µg/mL in coating buffer and incubated overnight at 4 °C. Using the previous ELISA protocol, plates were incubated with serial dilutions of samples to analyze, and bound VHH were further detected by HRP-conjugated Goat Anti-Alpaca IgG, VHH domain antibody. To estimate the EC50, four parameter logistic equation was fitted to resulting data. All indirect ELISA measurements were realized in triplicates. All washing steps were performed with Agilent BioTek washer dispenser (EL406 model). Unless stated otherwise, all reagents and equipment were purchased from ThermoFisher Scientific.

#### Microfluidic chips

Microfluidic droplet generator device and droplet analysis device were fabricated by classical soft lithography[4]. The droplet generator consisted of a 3 inlets flow focusing geometry with typical channel width of 20 µm, allowing water-in-oil emulsification of a co-flow of two aqueous phases. The droplet analyzer was composed of a 40 µm wide emulsion re-injection channel and two droplet spacing channels followed at the junction by a 30 µm long fluorescence detection region. For both designs a 42-43 µm thick layer of negative photoresist SU-8 2050 (KAYAKU Advanced Materials) was spin-coated on a silicon wafer following manufacturer's instructions and further exposed to UV light through a chromium transparency mask to obtain the positive master mold. Polydimethylsiloxane (PDMS, Sylgard 184 Silicone Elastomer, Dow Corning) was mixed with curing agent (ratio 10:1), poured onto the positive mold, thoroughly degassed, and cured for at least 1h at 65°C. The PDMS structure was peeled off the wafer and inlet/outlet holes were formed using biopsy punches (0.75 mm diameter, Robbins Instruments). The structure was then bonded to glass slide after oxygen-plasma activation. Microfluidic channels were treated with 1% Trichloro(1H,1H,2H,2H-perfluorooctyl)silane (Sigma-Aldrich) in Novec HFE-7500 fluorinated oil (3 M), flushed with HFE-7500, dried with compressed air and stored as is before experiments.

**Emulsion generation**

Polytetrafluoroethylene (PTFE) tubing with an inner diameter of 0.30 mm (Adtech, ThermoFisher Scientific) connected to low-retention pipette tips (ART™, ThermoFisher Scientific) was used to connect the droplet generator chip to gas-tight glass syringes (Hamilton). Flow rates were controlled by syringe pumps (Nemesys). Typically, 1400 µL/h for oil phase (HFE-7500 with 2 % of 008-Fluorosurfactant (RAN Biotechnologies)) and 400 µL/h for each aqueous phase, to obtain droplets of 30 pL at a production rate of 4000-5000 Hz. Prior to encapsulation, 300 nm diameter streptavidin-coated superparamagnetic beads (Bio-Adembeads Streptavidin, Ademtech) were coated according to manufacturer's protocol with HA Epitope Tag Antibody, Biotin conjugate (ThermoFisher Scientific) for NanoGFP expression or anti-c-Myc Antibody (Biolegend) in case of LaM-4 expression, at 60 µg of antibody per mg of beads. The functionalized beads dispersion was sonicated 2 min on ice to remove aggregates. Aqueous phase 1 typically contained antibody-coated magnetic beads at final concentration of 1.5 µg/µL, 40 % of PURExpress® A reagent, 30 pg/µl plasmidic DNA, RNAse inhibitor at 1.6 U/µL, 80 nM EGFP and nuclease-free water. In sequential droplet analysis with variable number of DNA copies (Fig. 4C) all emulsions were pooled and analyzed simultaneously using fluorescence barcoding by addition of 75 to 530 nM Sulforhodamine B (Sigma-Aldrich) in aqueous phase 1. In specificity assays (Fig. 5), aqueous phase 1 also contained 160 nM mCherry (OriGene). For the EGFP detection by a secondary antibody (Fig. S10) we used in phase 1 an anti-GFP IgG Alexa647 conjugate (Sigma-Aldrich) at 80 nM. Aqueous phase 2 always contained 40% of PURExpress® A reagent and 60% of PURExpress® B reagent. The emulsion was incubated for further measurements in a 0.5 mL Eppendorf-type tube filled with HFE-7500 containing 0.5% 008-Fluorosurfactant and closed with a home-made inlet-outlet PDMS connector for further reinjection of the emulsion.

**Laser-assisted measurements of droplet fluorescent profile**

The collector tube was then inserted into a ring-shaped neodymium magnet (Magnet-shop) and placed in a 37 °C incubator for 3 h. To prevent evaporation inlet tubing was closed, and outlet tubing was connected to a syringe containing the same oil phase as collector tube. After incubation, droplets were reinjected using syringe pumps into a droplet analysis chip placed between flat neodymium magnets. Measurements of droplet fluorescent profiles were performed at 500 Hz on a microfluidic platform similar to that described in ref [5] using a multicolor laser source L6Cc (Oxxius) for excitation and bandpass filters (Semrock) combined with photomultiplier tubes (H10723 series, Hamamatsu) for collection of emitted photons. Laser beam was line-shaped and oriented parallel to the capture scaffold to maximize specific signal. EGFP signal was obtained using the 488 nm laser and 525/40 emission filter (PMT2) while Sulforhodamine B and mCherry signal were measured using 561 nm laser and 593/46 filter (PMT3). Finally, the signal of Anti-GFP IgG Alexa647 conjugate was assessed with 638 nm laser and 708/75 filter (PMT4). Maximal fluorescence intensities detected within single droplets were followed and recorded by a custom LabView software and data analysis carried out in FlowJo.

##### Fluorescence Microscopy Imaging

Custom made 50 µm deep microfluidic glass chambers (IMT Masken und Teilungen) equipped with inlet and outlet valves were pre-filled with HFE-7500, loaded with 80 µl of emulsion to image and placed between magnets for the time of imaging. Incubation at 37°C was carried out on Heatable Universal Mounting Frame (A-H R S, PeCon) controlled by TempModule S (Zeiss). Fluorescence imaging was realized by inverted epifluorescence microscope AxioObserver Z1 (Zeiss) equipped with EC Plan-Neofluar 10x/0.30 objective, large spectrum mercury arc lamp (HBO100, Zeiss) and GFP filter set (Ex: 475/40, Dc: 500, Em: 530/50, Zeiss). An sCMOS camera (ORCA-Flash4.0, Hamamatsu) controlled by ZEN lite software (Zeiss) was used for brightfield and fluorescence images acquisition. Images were treated using a home-made program in Fiji macro. Briefly, brightfield images were manually thresholded to detect the capture scaffold or drop background by gray-level segmentation. Particle analysis plugin was then used to measure the mean pixel fluorescence from detected regions. A minimum of 300 objects were analyzed per condition (Fig. 3C).

##### Plasmid map

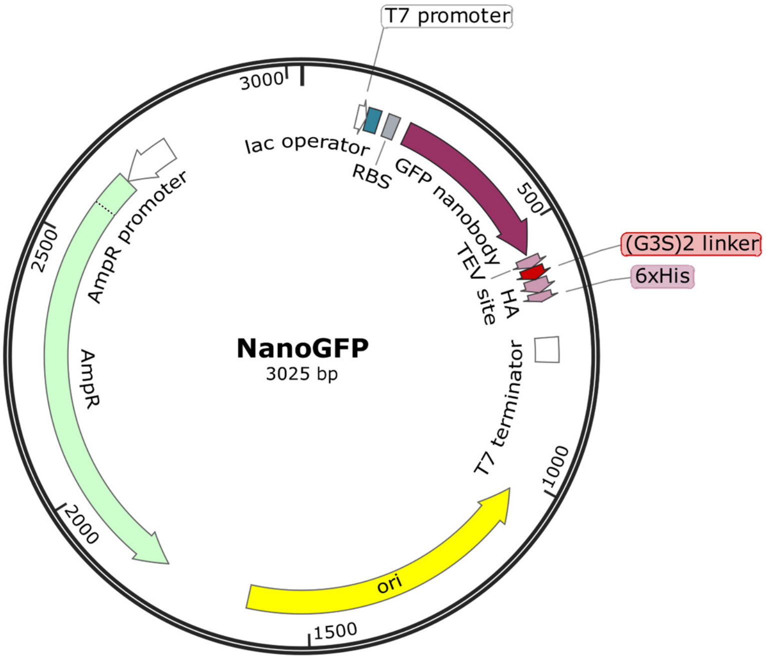


Primers used for subcloning of antibody genes into pRSET-B vector, MM1 (forward) and MM2 (reverse) were obtained from Sigma-Aldrich.

Subcloning primers:

| Primer | Sequence |
|---|---|
| MM1 | GGTATATCTCCTTCTTAAAGTTA |
| MM2 | ACTAGTAGGCTGCTAACAAAGCCCGA |

List of features with corresponding sequences:

| Feature | Sequence |
|---|---|
| T7 promoter | TAATACGACTCACTATAGG |
| T7 terminator | CTAGCATAACCCCTTGGGGCCTCTAAACGGGTCTTGAGGGG TTTTTTG |
| Lac operator (lacO) | GGAATTGTGAGCGGATAACAATTCC |
| Ribosome binding site (RBS) | TTTGTTTAACTTTAAGAAGGAGA |
| TEV site | GAGAACCTCTACTTCCAATCG |
| (G3S)2 linker | GGTGGCGGTAGTGGTGGCGGTAGT |
| Hemagglutinin tag (HA), only anti-GFP antibodies | TATCCGTATGATGTTCCGGATTATGCA |
| c-Myc tag, only anti-mCherry antibody | GAACAAAAACTCATCTCAGAAGAGGATCTG |
| Histidine tag (6xHis) | CACCACCATCATCACCAT |
| NanoGFP | ATG GCA CAG GTG CAG CTG GTT GAA AGC GGT GGT GCA CTG GTT CAG CCT GGT GGT AGC CTG CGT CTG AGC TGT G CA GCA AGC GGT TTT CCG GTT AAT CGT TAT AGC ATG CG T TGG TAT CGT CAG GCA CCG GGT AAA GAA CGT GAA TG GGTT GCA GGT ATG AGC AGT GCC GGT GAT CGT AGC AG C TAT GAA GAT AGC GTT AAA GGT CGT TTT ACC ATC AG C CGT GAT GAT GCA CGT AAT ACC GTT TAT CTG CAA ATG AAT AGC CTG AAA CCG GAA GAT ACC GCA GTG TAT TAT TGC AAT GTT AAC GTG GGC TTT GAA TAT TGG GGT CAG GGC ACC CAG GTT ACC GTT AGC AGC |
| LaG-2 | ATG CAG GTT CAG CTG GTT GAA AGC GGT GGT GGT CTG GTT CAG GCA GGT GGT AGC CTG CGT CTG AGC TGT GCA GCA AGC GGT CGC ACC TTT AGC AAC TAT GCG ATG GGC TGG TTT CGT CAG GCA CCG GGT AAA GAA CGT GAA TTT GTT GCA GCA ATC AGC TGG ACC GGT GTT AGC ACC TAT TAT GCG GAT AGC GTT AAA GGT CGT TTT ACC ATC AGC C GT GAT AAC GAT AAA AAT ACC GTT TAT GTG CAA ATG A AT AGC CTG ATC CCG GAA GAT ACC GCA ATC TAT TAT T GC GCG GCG GTG CGC GCG CGC AGC TTT AGC GAT ACC T |

| | |
|---|---|
| | AT AGC CGC GTT AAC GAA TAT GAT TAT TGG GGT CAG G GC ACC CAG GTT ACC GTT |
| LaG-6 | ATG CAG GTT CAG CTG GTT GAA AGC GGT GGT GGT CTG GTT CAG GCA GGT GGT AGC CTG CGT CTG AGC TGT GCA GCA AGC GGT CGC ACC TTT AGC ACC AGC GCG ATG GCG TGG TTT CGT CAG GCA CCG GGT AAA GAA CGT GAA TTT GCA GCA GGT ATT ACC TGG ATT AGC AGC AGC ACC TAT TAT ACC GAT AGC GTT AAA GGT CGT TTT ACC ATC AGC C GT GAT AAT GCA AAA AAT ACC GTT TAT CTG CAA ATG A AT AGC CTG AAA CCG GAA GAT ACC GCA GTG TAT TAT T GC GCG GCG AAA AGC GAA GGC TAT TTT GGC TTT CCG C GC GTG GAA AAC GAA TAT CCG TAT TGG GGT CAG GGC A CC CAG GTT ACC GTT |
| LaG-18 | ATG GCG CAG GTT CAG CTG GTT GAA AGC GGT GGT GGT CTG GTT CAG ACC GGT GGT AGC CTG AAA CTG AGC TGT ACC GCA AGC GTT CGC ACC CTG AGC TAT TAT CAT GTG GGC TGG TTT CGT CAG GCA CCG GGT AAA GAA CGT GAA TTT GTT GCA GGC ATT CAT CGC AGC GGC GAA AGC ACC TTT TAT GCC GAT AGC GTT AAA GGT CGT TTT ACC ATC A GC CGT GAT AAT GCA AAA AAT ACC GTT CAT CTG CAA A TG AAT AGC CTG AAA CCG GAA GAT ACC GCA GTG TAT T AT TGC GCG CAG CGC GTG CGC GGC TTT TTT GGC CCG CT G CGC AGC ACC CCG AGC TGG TAT GAT TAT TGG GGT CA G GGC ACC CAG GTT ACC GTT AGC |
| LaM-4 | ATG GCG CAG GTT CAG CTG GTT GAA AGC GGT GGT AGC CTG GTT CAG CCT GGT GGT AGC CTG CGT CTG AGC TGT G CA GCA AGC GGT CGC TTT GCG GAA AGC AGC AGC ATG G GC TGG TTT CGT CAG GCA CCG GGT AAA GAA CGT GAA T TTGTT GCA GCG ATT AGC TGG AGC GGC GGC GCG ACC A AC TAT GCG GAT AGC GCA AAA GGT CGT TTT ACC CTG A GC CGT GAT AAT ACC AAA AAT ACC GTT TAT CTG CAA A TG AAT AGC CTG AAA CCG GAT GAT ACC GCA GTG TAT T AT TGC GCG GCG AAC CTG GGC AAC TAT ATT AGC AGC A AC CAG CGC CTG TAT GGCTAT TGG GGT CAG GGC ACC C AG GTT ACC GTT AGC AGC |
| YFAST | ATG GAG CAT GTT GCC TTT GGC AGT GAG GAC ATC GAG AAC ACT CTG GCC AAA ATG GAC GAC GGA CAA CTG GAT GGG TTG GCC TTT GGC GCA ATT CAG CTC GAT GGT GAC GGG AAT ATC CTG CAG TAC AAT GCT GCT GAA GGA GAC ATC ACA GGC AGA GAT CCC AAA CAG GTG ATT GGG AAG AAC TTC TTC AAG GAT GTT GCA CCT GGA ACG GAT TCT CCC GAG TTT TAC GGC AAA TTC AAG GAA GGC GTA GCG TCA GGG AAT CTG AAC ACC ATG TTC GAA TGG ATG ATA CCG ACA AGC AGG GGA CCA ACC AAG GTC AAG GTG CAC ATG AAG AAA GCC CTT TCC GGT GAC AGC TAT TGG GTC TTT GTG AAA CGG GTG |

#### 2. Supplementary Figures S1-S10

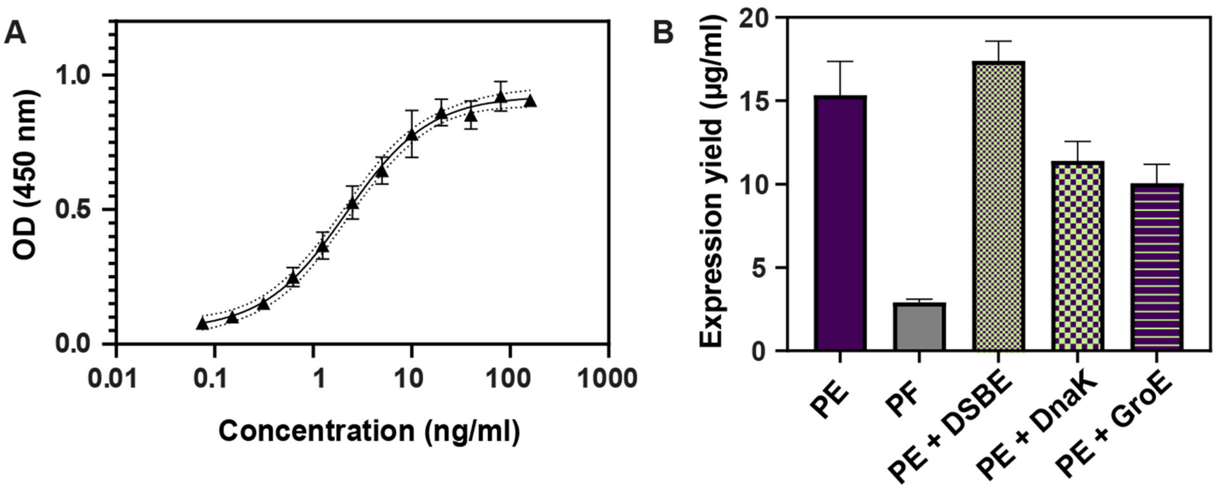


**Figure S1. Measurement of NanoGFP expression yield in bulk.** A) Standard curve obtained by sandwich ELISA with dilutions of commercial anti-GFP VHH (ChromoTek). Dotted lines correspond to the asymptotic 95 % confidence interval. B) Expression yield of NanoGFP synthesized (3 h, 37 °C) with PURExpress® (PE), PURE*frex*® (PF), PE supplemented before incubation with disulfide bond enhancer (DSBE, NEB), with DnaK mix of chaperones (GeneFrontier) and GroE chaperone (Genefrontier), (means ± standard deviation, *n* = 3). All supplements were added to expression medium as per manufacturer's instructions.

**A**

| Clone | Size (kDa) | Nature of CDR3 | $K_D$ (theoretical) | Homology with NanoGFP |
|---|---|---|---|---|
| NanoGFP | 16.4 | Concave (6 AA) | 0.23 / 0.32 / 1 nM | - |
| LaG-2 | 17.4 | Convex (18 AA) | 16 nM | 65.1% |
| LaG-6 | 17.2 | Convex (17 AA) | 0.3 µM | 66.1% |
| LaG-18 | 17.7 | Convex (18 AA) | 3.8 µM | 66.4% |

**B**

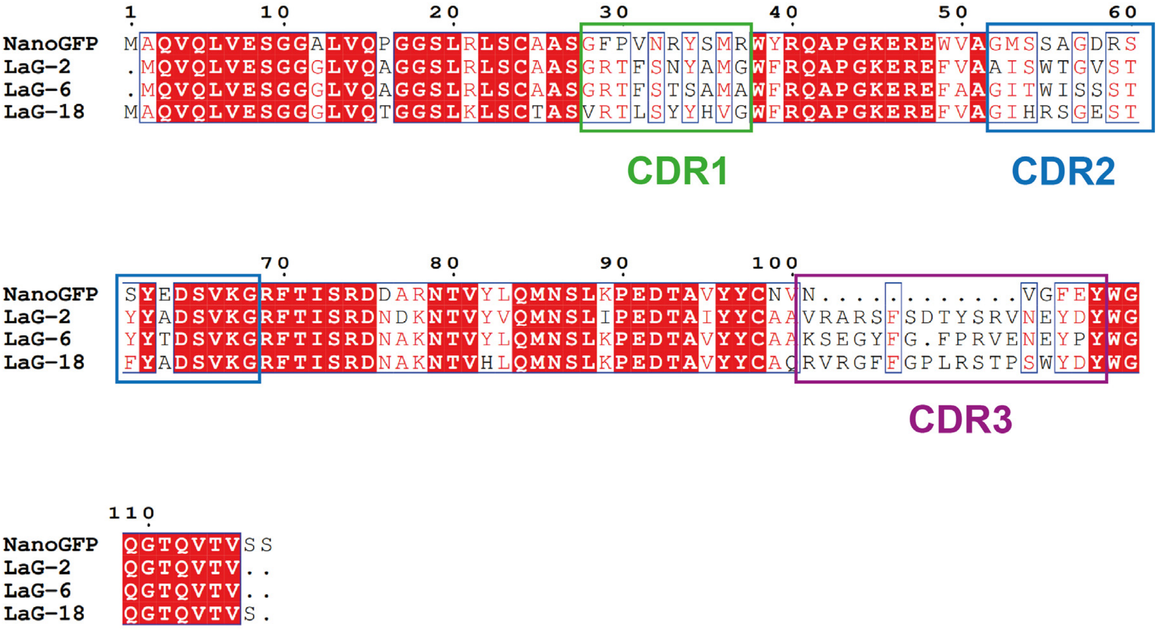


**Figure S2. Properties of cell-free expressed anti-GFP VHH sequences.** A) Comparative table of VHH characteristics. The indicated size comprises features located downstream VHH sequence (e.g., linker, HA epitope tag and 6xHistidine tag). Theoretical $K_D$ of NanoGFP corresponds to previously published results obtained by different experimental methods.[2,6,7] Indicated sequence homology is the one of VHH translated sequences only, downstream features were excluded. B) Sequence alignment generated by ESPript 3.0.[8] Amino acids both framed and red highlighted display identity between all sequences, framed-only amino acids show partial sequence identity.

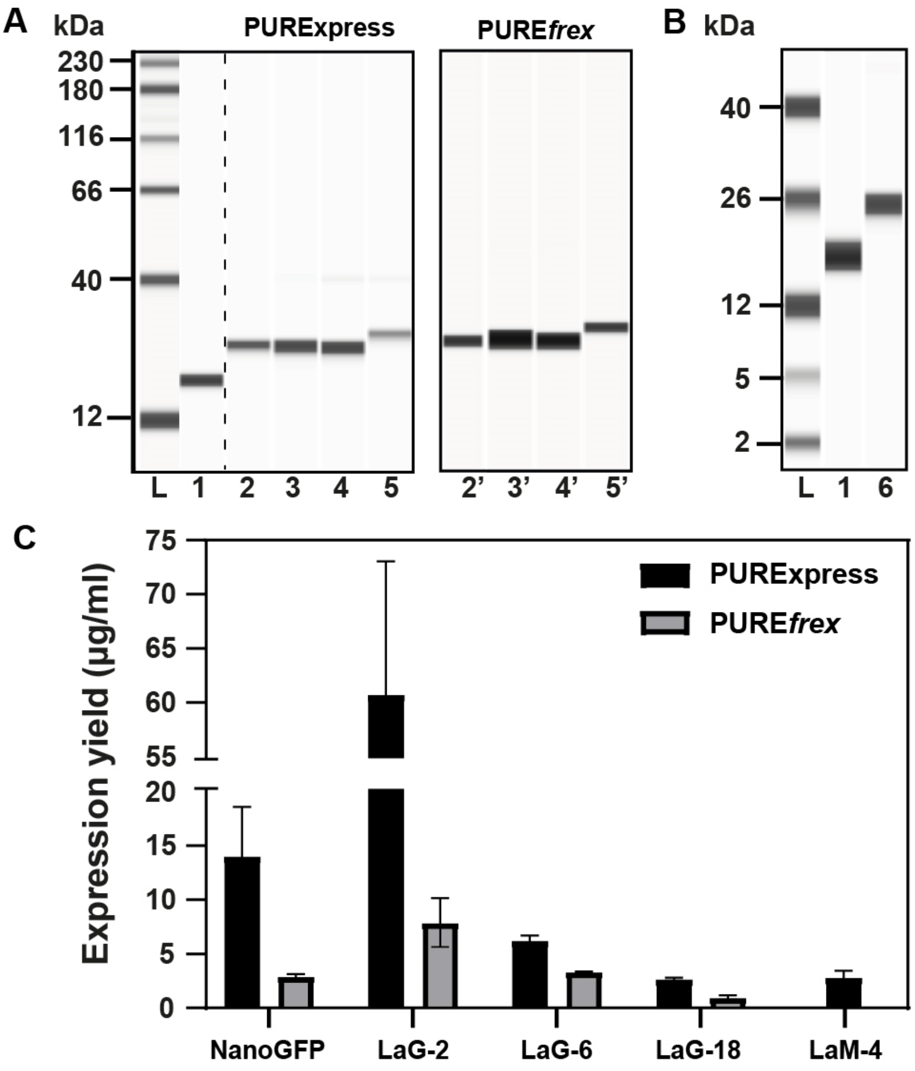


**Figure S3. Characterization of 4 anti-GFP VHHs, expressed in bulk by PURExpress® and PUREfrex® and anti-mCherry LaM-4 expressed PURExpress®.** A) Capillary western blot analysis of anti-GFP VHHs. Electrophoretogram displays the molecular weight ladder (lane L), commercially available anti-GFP VHH (lane 1), NanoGFP (lane 2), LaG-2 (lane 3), LaG-6 (lane 4) and LaG-18 (lane 5). VHH synthetized in PURExpress®: left, in PUREfrex®: right. B) Capillary western blot analysis of commercially available anti-GFP VHH (lane 1) and LaM-4 (lane 6). C) Expression yield measured by ELISA titration (calibration curve in Fig.S1) after 3 h of expression at 37 °C (means ± standard deviation, *n* = 3).

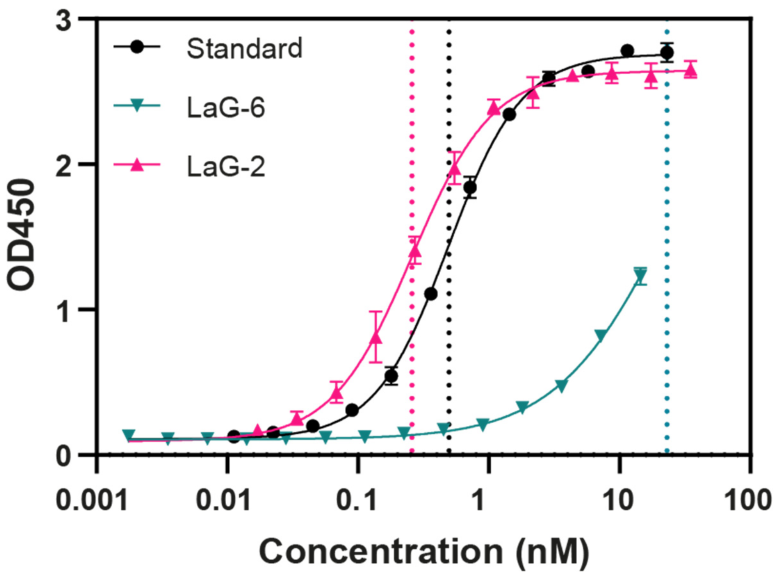


**Figure S4. Dose-response curves of commercially available anti-GFP VHH (Standard), cell-free expressed LaG-2 and cell-free expressed LaG-6.** Dotted lines indicate the apparent dissociation constant, for the standard: $K_D^{app}$ = 0.49 ± 0.03 nM, for the LaG-2: $K_D^{app}$ = 0.26 ± 0.02 nM and for the LaG-6: $K_D^{app}$ = 19.92 ± 1.35 nM. LaG-2 and LaG-6 were syntheiszed in PURExpress® at [DNA] = 4 ng/µL, 37 °C, 3 h. The affinity LaG-18 being too low (theoretical $K_D$ = 3.8 µM), it could not be assessed by indirect ELISA.

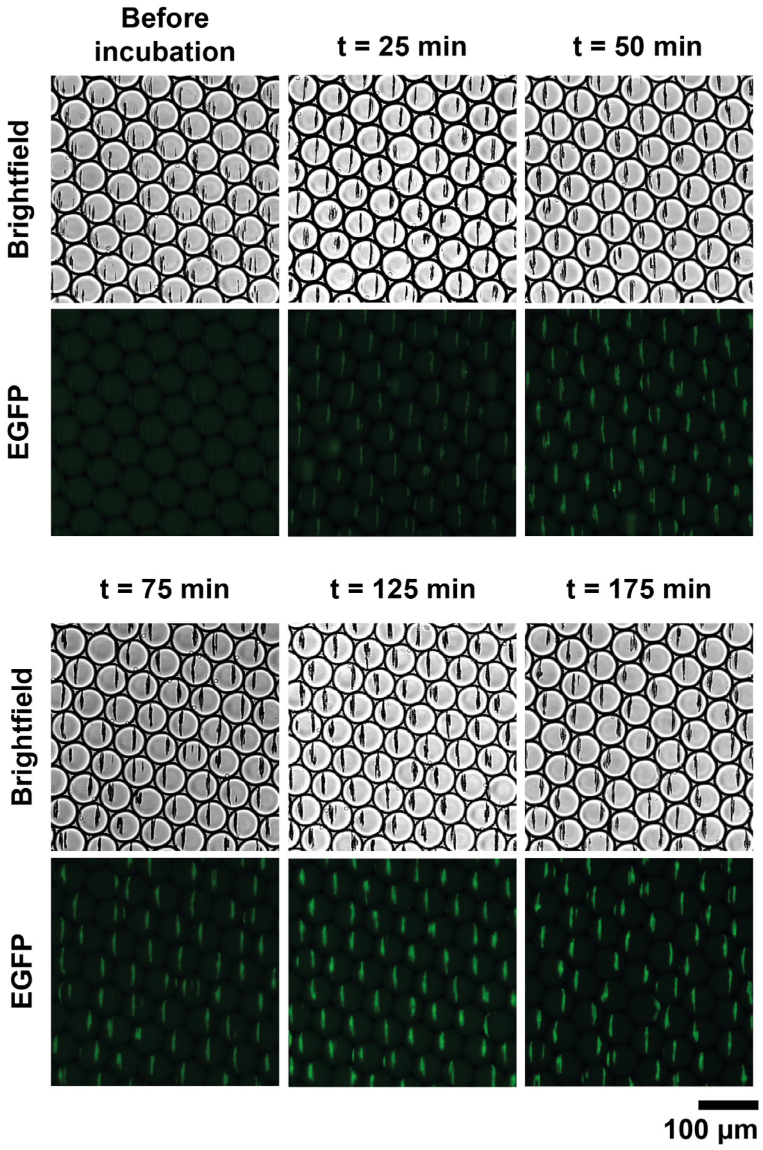


**Figure S5. Monitoring of NanoGFP expression and EGFP binding by fluorescence microscopy.** Directly after encapsulation, the emulsion was loaded into a glass microfluidic chamber and imaged before and during incubation at 37 °C. To prevent photobleaching of EGFP, different regions of chamber were imaged each time.

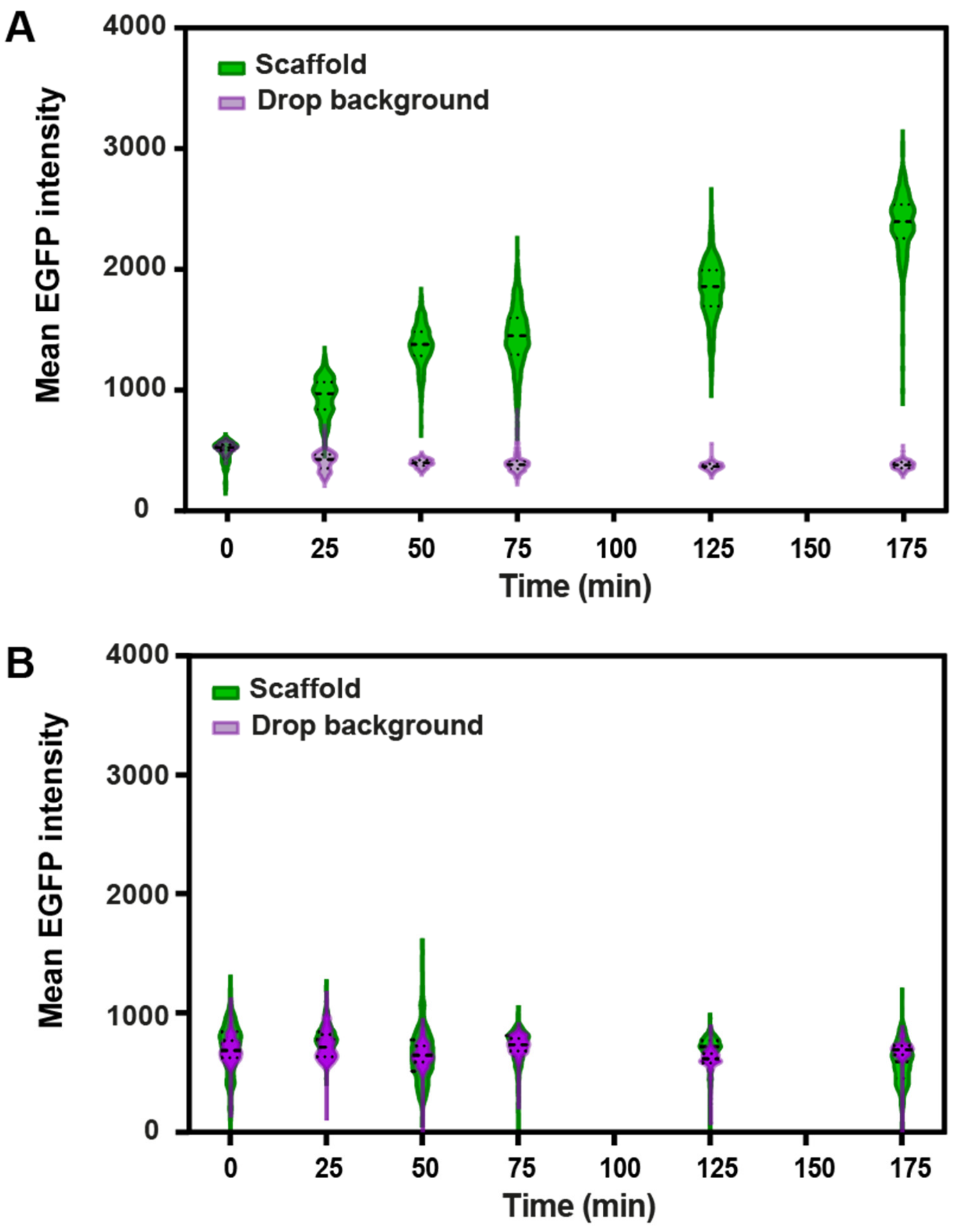


**Figure S6. Violin plot of the temporal evolution of mean EGFP intensity measured on the capture scaffold and in the drop background with (A) and without (B) DNA coding for NanoGFP.** Upon encapsulation, the emulsion was loaded into a microfluidic glass chamber, incubated at 37 °C and imaged at different time-points. Mean EGFP intensity of 300 droplets was assessed by particle analysis on background-subtracted images (dashed line: median, dotted lines: lower and upper quartiles).

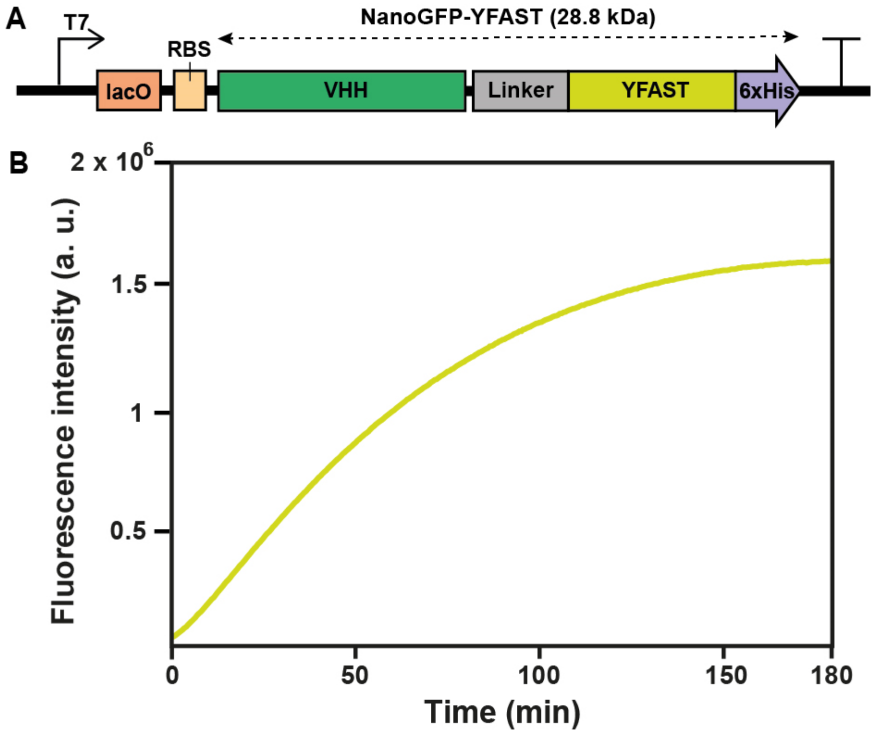


**Figure S7. Bulk expression of NanoGFP fused to a fluorogenic YFAST[9] reporter.** A) Schematic representation of the DNA template used to follow NanoGFP expression temporally. The HA tag was replaced by YFAST protein. B) Fluorescence intensity of NanoGFP-YFAST expressed with PURExpress® at [DNA] = 4 ng/µL with 50 µM HMBR was monitored in a 10 µL reaction by qPCR (QuantStudio 4, ThermoFisher) with 470/15 nm excitation and 558/11 nm emission channel over 3 h.

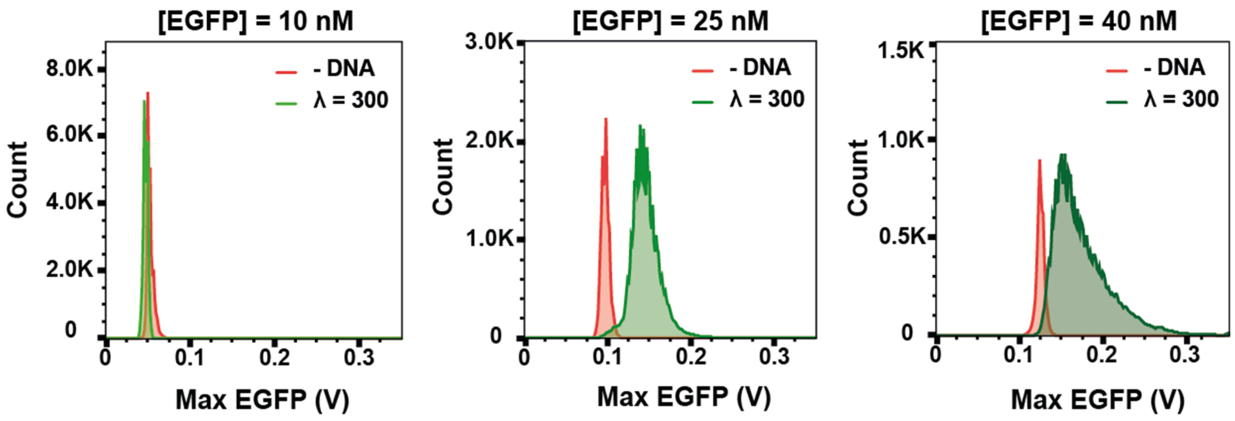


**Figure S8. NanoGFP expression with varying EGFP concentration.** Emulsions containing 10, 25 and 40 nM EGFP were compared to the condition without DNA. All emulsions contained for λ = 300 plasmids per drop and were incubated at 37 °C for 3 h.

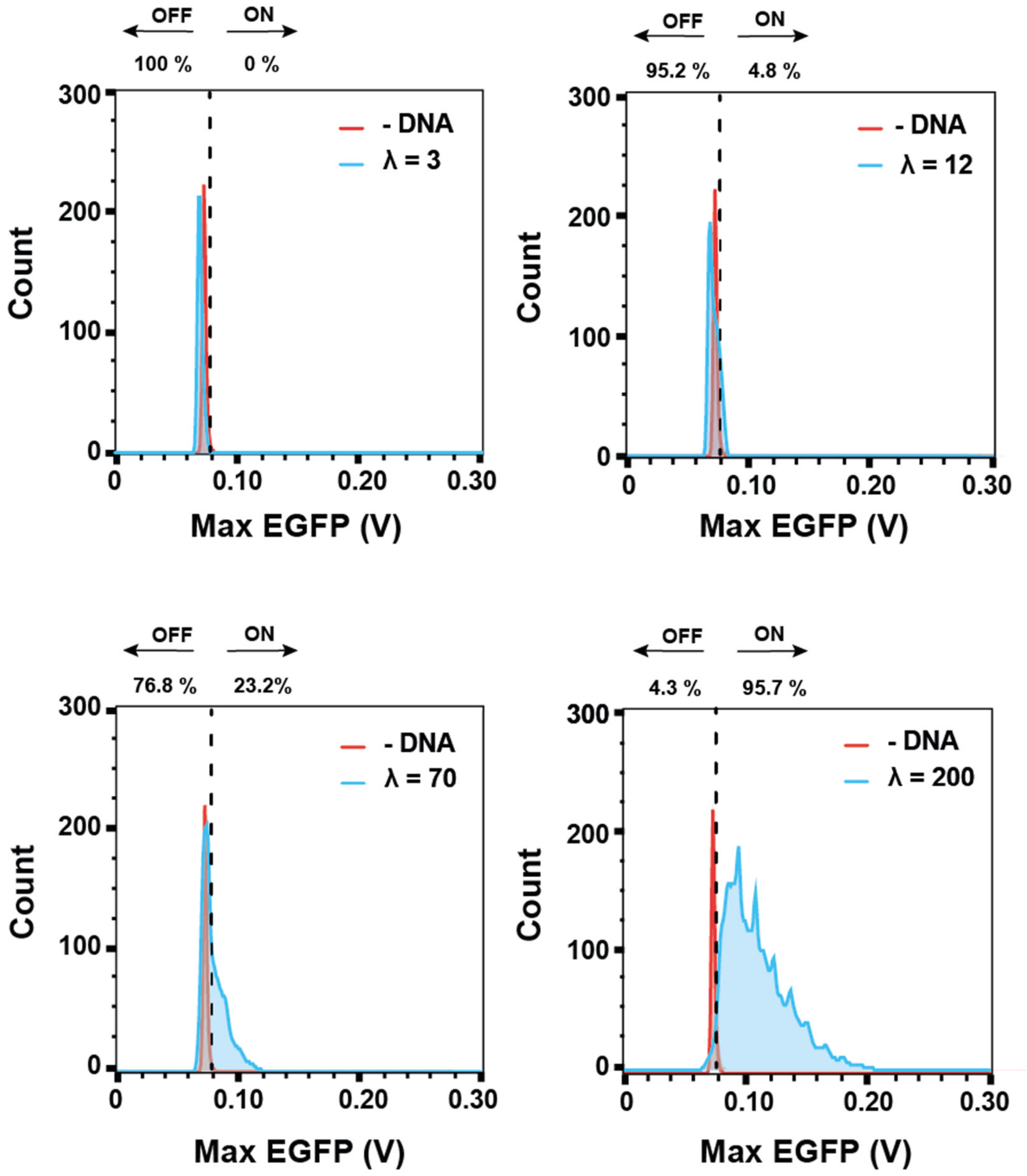


**Figure S9. NanoGFP expression assessed by sequential analysis for λ = 3, 12, 70 and 200 plasmids per drop.** Each condition was compared to the emulsion without coding DNA (-DNA). The dashed line indicates the threshold above which droplets are defined as positive. The proportion of positive (ON) and negative (OFF) droplets is indicated above each histogram. All emulsions contained 40 nM of EGFP and were incubated at 37 °C for 3 h.

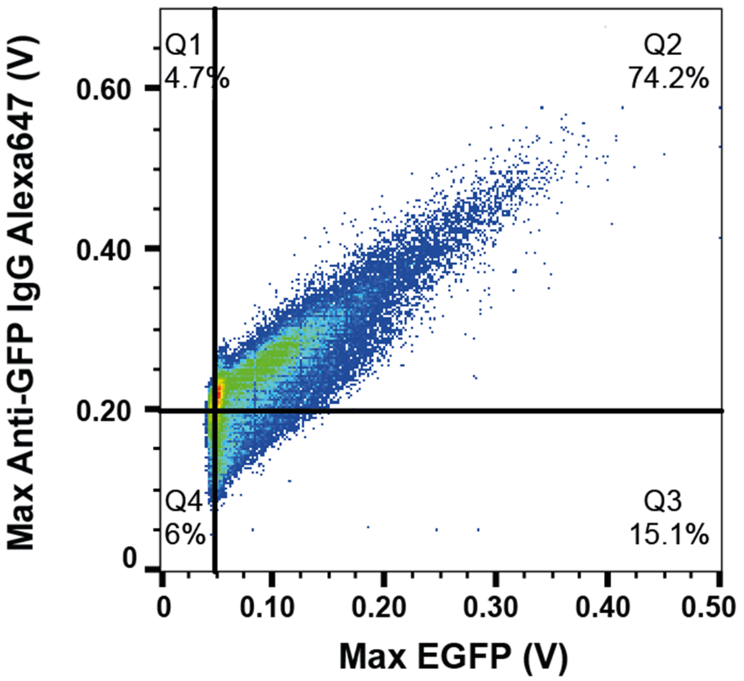


**Figure S10. Scatter plot of NanoGFP expression with secondary antibody for detection of EGFP.** The thresholds separating the plot into Q1-4 represent the highest Max Anti-GFP IgG Alexa647 and Max EGFP detected in condition without DNA. Q1 + Q2 indicate droplets positive in the red channel (anti-GFP IgG Alexa647 signal) and Q2 + Q3 those positive in the green channel (EGFP signal). Q4 represents droplets that remained negative. The emulsion contained $\lambda = 300$ plasmids per drop and 40 nM of both EGFP and anti-GFP IgG Alexa647. It was analyzed after 3 h of incubation at 37°C ($N = 83797$).

#### 3. Supplementary Table S1

**Table S1.** The list of described sequentially analyzed immunoassay conditions with the total number of droplets detected ($N_{det}$) and represented as histograms ($N_{plot}$).

| Figure | λ | Channel | $N_{det}$ | $N_{plot}$ |
|---|---|---|---|---|
| 4B, S8 | 0 | EGFP | 42378 | 6334 |
| 4B, S8 | 300 (NanoGFP) | EGFP | 37590 | 37590 |
| 4C | 0 | EGFP | 24107 | 782 |
| 4C | 3 (NanoGFP) | EGFP | 24894 | 1021 |
| 4C | 12 (NanoGFP) | EGFP | 6840 | 1681 |
| 4C | 70 (NanoGFP) | EGFP | 42001 | 3003 |
| 4C | 200 (NanoGFP) | EGFP | 8113 | 8113 |
| 5 | 0 | EGFP | 40865 | 9802 |
| 5 | 0 | mCherry | 77263 | 8005 |
| 5 | 300 (NanoGFP) | EGFP | 31086 | 31086 |
| 5 | 300 (NanoGFP) | mCherry | 31086 | 9104 |
| 5 | 300 (LaM-4) | EGFP | 27588 | 9634 |
| 5 | 300 (LaM-4) | mCherry | 27588 | 27588 |
| S8 | 0 | EGFP (10 nM) | 32313 | 32313 |
| S8 | 300 (NanoGFP) | EGFP (10 nM) | 33247 | 33247 |
| S8 | 0 | EGFP (25 nM) | 43702 | 17068 |
| S8 | 300 (NanoGFP) | EGFP (25 nM) | 49818 | 49818 |

### 4. Supplementary Text S1

**Text S1.**

The characteristic time for Brownian diffusion in 3D of a molecule (here NanoGFP:EGFP complex, 44.4 kDa) is given by (1), where diffusion coefficient D is defined by Stokes-Einstein relation (2). To estimate the characteristic diffusion time of 4.5 s in our system we used the following parameters: $L$ = 40 µm (drop diameter), $T$ = 310.15 K, $\eta$ = $1.3\cdot10^{-3}$ $N.s.m^{-2}$ (viscosity of 20 % glycerol solution at 310.15 K), $r$ = 3 nm (radius of 44.4 kDa protein).

(1) $$\tau = \frac{L^2}{6D}$$

(2) $$D = \frac{k_B T}{6\pi\eta r}$$

$L$ : diffusion length

$k_B$ : Boltzmann constant

$T$ : temperature

$\eta$ : viscosity

$r$ : molecule radius
